# A scalable screening approach reveals phylogenetic patterns of bacterial antagonism against potato pathogens

**DOI:** 10.64898/2026.09.22.753498

**Authors:** Vivien Pichon, Alisson Gillon, Gaëlle Ruffieux, Tom Luethi, Camila Morales Undurraga, Mout De Vrieze, Floriane L’Haridon, Ola Abdelrahman, Laure Weisskopf

## Abstract

Microbial biological control agents (mBCAs) offer a promising alternative to conventional disease management, but identifying effective strains requires screening large microbial collections against many pathogens. Here, we developed a scalable screening approach to characterize the antagonistic activity of potato-associated bacteria against multiple pathogens and investigated how bacterial origin and phylogeny relate to antagonistic phenotypes. A collection of 600 bacterial strains isolated from two potato cultivars, three plant compartments using four cultivation media was screened against six major potato pathogens: the bacteria *Dickeya solani* and *Pectobacterium carotovorum*, the fungi *Alternaria solani* and *Rhizoctonia solani*, and the oomycetes *Phytophthora infestans* and *Pythium ultimum*. A scalable confrontation method was developed for non-filamentous bacteria, while complementary lower-throughput assays were used for filamentous strains. This approach enabled successful assessment of 92% of non-filamentous and 97% of filamentous strain-pathogen combinations. Isolation cultivar, plant compartment and cultivation medium had only limited effects on the proportion of antagonistic strains. In contrast, pathogen sensitivity varied markedly: *P. infestans* was the most susceptible pathogen, whereas *D. solani* and *P. carotovorum* were generally resistant to bacterial antagonism. Antagonistic activity clustered phylogenetically, with *Bacillus* and *Streptomyces* containing many broad-spectrum inhibitors. In addition, several taxa showed preferential activity against the oomycetes, including *P. infestans*-specific antagonists from the *Pseudomonas* genus and *P. ultimum*-specific antagonists from the *Frigoribacterium*, *Curtobacterium*, *Pedobacter* and *Phyllobacterium* genera. Overall, multi-pathogen screening revealed distinct generalist and specialist antagonistic profiles and identified candidate strains and taxa for further evaluation as mBCAs.

**Highlights:**

- A scalable confrontation method enabled the screening of 400 non-filamentous bacterial strains against six potato pathogens. Two hundred filamentous strains from the Actinomycetes class required separate lower-throughput methods because growth conditions were incompatible with the scalable assay.
- The screening of these 600 potato-associated bacterial strains revealed distinct phylogenetic patterns of broad-spectrum and pathogen-specific antagonism across six major potato pathogens.
- Pathogen sensitivity differed strongly among taxonomic groups: the bacterial pathogens *Dickeya solani* and *Pectobacterium carotovorum* were the most resistant, the oomycetes *Phytophthora infestans* and *Pythium ultimum* were the most sensitive, and the fungi *Alternaria solani* and *Rhizoctonia solani* showed intermediate sensitivity.
- Specialist antagonists were identified in several bacterial taxa, including Micrococcales, Sphingobacteriales and Rhizobiales against *P. ultimum*, and Streptomycetales and Pseudomonadales against *Phytophthora infestans*.
- The bacterial and fungal pathogens were almost exclusively inhibited by broad-spectrum bacterial strains from the *Streptomyces* and *Bacillus* genera.

## Introduction

Potato (*Solanum tuberosum* L.) is one of the most important staple crops worldwide and plays a crucial role in global food security (FAO, 2023). Its adaptability to diverse environmental conditions and high productivity have contributed to its widespread production worldwide. However, its cultivation remains highly challenging due to severe yield and quality losses caused by plant diseases (Kumar Tiwari et al., 2021; Shi et al., 2023).

Potato crops are susceptible to numerous pathogens belonging to distinct taxonomic groups, including fungi, oomycetes, bacteria, and viruses (Chiarini et al., 2026). These pathogens colonize various plant compartments and employ distinct infection strategies, complicating effective disease management. Among the most important potato diseases are late blight caused by the oomycete pathogen *Phytophthora infestans;* early blight caused by the fungus *Alternaria solani*; stem canker and black scurf caused by the fungus *Rhizoctonia solani; root rot and tuber decay caused by the oomycete Pythium ultimum*; blackleg caused by the bacteria *Dickeya solani* and soft rot disease caused by the bacteria *Pectobacterium carotovorum*. Together, they reduce crop productivity, undermine tuber quality, and generate substantial economic losses throughout potato production and storage (Czajkowski et al., 2011; Fry, 2008; Mugao, 2023; Parveen and Sharma, 2015; Tsror, 2010).

Current disease management relies primarily on integrated approaches that combine resistant cultivars, cultural practices, and, most importantly, chemical pesticides (Kirk et al., 2005; Nærstad et al., 2007; Sharma et al., 2015). Although chemical control remains an important tool for potato production, its long-term sustainability is increasingly questioned. Intensive and repeated fungicide applications have contributed to the emergence of resistant pathogen populations, reducing treatment efficacy, and leading to an increasing need for alternative management strategies (Matson et al., 2015). In addition, growing concerns regarding the environmental and sanitary costs of conventional treatments drive the progressive ban of several active compounds, pressing the need to find substitute solutions (Zhou et al., 2021).

Consequently, microbial biological control agents (mBCAs) have attracted increasing attention as potential sustainable alternatives to suppress plant pathogens while reducing agricultural reliance on synthetic pesticides (De Pessemier et al., 2026; Marchand, 2023; Shi et al., 2023; Villavicencio-Vásquez et al., 2025).

The plant microbiota represents a particularly promising reservoir of potential mBCAs, as these microbes are naturally able to colonize and interact with the plants. In addition, the rhizosphere soil, roots, and leaves harbor diverse bacterial communities, including several genera such as *Bacillus*, *Pseudomonas*, *Streptomyces* and *Paenibacillus* that have been investigated for their antagonistic activity against potato pathogens (Brewer and Larkin, 2005; Elliott et al., 2009; González-Franco and Robles-Hernández, 2009; Shi et al., 2023).

However, taxonomic characterization of microbial community composition alone does not identify which individual components actively contribute to pathogen suppression, demonstrating the importance of the functional characterization of cultivated microbial isolates. Cultivable microbes only represent a small fraction of the total communities, although this proportion increases with plant vicinity (Buyer et al., 2002; Youseif et al., 2021). Additionally, antagonistic microorganisms usually represent only a subset of these cultivable microbes, making the screening of large numbers of individual isolates a necessity for the identification of promising biological control candidates (Köhl et al., 2011).

Traditionally, microbial antagonism has been assessed *in vitro* using confrontation assays in which individual isolates are evaluated against a single pathogen under controlled conditions. While such assays can successfully identify the antagonistic potential of single isolates, they are labor-intensive, time-consuming, and may be difficult to standardize when applied to large microbial collections due to the diverse growth and physiological properties of their members (Köhl et al., 2011). Besides, evaluating isolates against a single pathogen provides limited information about their overall biocontrol potential. Thus, testing the same collection against multiple pathogens substantially increases the amount of useful information, revealing whether a strain exhibits broad-spectrum activity or is specialized against one or few pathogens (Köhl et al., 2019; Raymaekers et al., 2020). The characterization of the antagonistic spectrum is relevant for mBCA development, as isolates with broad-spectrum activity may offer protection of plants against multiple pathogens, but are also more likely to affect non-target microorganisms, in contrast to isolates with a narrower range of activity (Köhl et al., 2019; Price-Christenson and Yannarell, 2023). However, expanding screenings to multiple pathogens also increases methodological complexity, as the assays must accommodate pathogens’ growth conditions, which often vary from one pathogen to another. Consequently, the development of reliable, reproducible, and scalable screening approaches is essential for accelerating the discovery of promising mBCAs.

In this study, we took advantage of a recently assembled collection of potato-associated bacteria (Pichon et al., 2026) to develop a high-throughput testing protocol fitting multiple pathogens from different taxonomic groups, while investigating the following scientific questions: i) Do isolation characteristics (cultivar, phyllosphere vs. rhizosphere, cultivation medium) influence the proportion of retrieved antagonistic strains?; ii) Do pathogens belonging to the same taxonomic group (fungi, oomycete, bacteria) show similar profiles of sensitivity to the tested isolates?; iii) Are certain bacterial taxa richer in antagonistic strains against specific pathogens? If yes, which taxa are they?

The bacterial collection we used in this study comprised 600 strains belonging to 62 different genera, including filamentous (from the Actinomycetes class) and non-filamentous isolates. They were screened against six pathogens representing the major taxonomic groups affecting potato: the oomycetes *Phytophthora infestans* and *Pythium ultimum*, the fungi *Alternaria solani* and *Rhizoctonia solani*, and the bacteria *Dickeya solani* and *Pectobacterium carotovorum*. By evaluating the same bacterial collection against multiple pathogens, we sought i) to identify isolates displaying either broad-spectrum or pathogen-specific antagonistic activity, ii) to compare the sensitivity of each pathogen to inhibition by different bacteria, and iii) to assess the suitability of our screening methods for large-scale functional characterization.

## Material and methods

### Strains and cultivation media

#### Collection strains

The detailed description of the isolation protocol of the strain collection can be found in Pichon et al. 2026. Briefly, microbial and fungal strains were isolated from the rhizosphere, root and phyllosphere of potato plants of the Bintje and Innovator cultivars. Plant-associated samples were finely ground and plated onto four different culture media supplemented with selective antibiotics to inhibit unwanted microbial growth. Media were supplemented either with nystatin (10 μg/mL) to inhibit fungal growth or with ampicillin and kanamycin (250 μg/mL and 25 μg/mL, respectively) to inhibit bacterial growth.

The growth media used for isolation were selected to maximize the diversity of isolated microorganisms. Ten percent Tryptic Soy Agar (TSA, Sigma-Aldrich, Darmstadt, Germany) with nystatin was used as a nutrient-rich medium providing abundant carbon and nitrogen sources. Twenty percent Potato Dextrose Agar (PDA; Carl Roth, Karlsruhe, Germany), supplemented with ampicillin and kanamycin was primarily used to isolate fungi, but also included bacterial isolates. Actinomycete Isolation Agar (ActIA; Sigma-Aldrich), supplemented with nystatin, was used to favour the growth of actinomycetes. Finally, Artificial Root Exudate medium (ARE) (Schmidt et al., 2016b) was supplemented either with ampicillin or with nystatin and kanamycin. This nutrient-poor medium with organic acids was included to promote the growth of rhizosphere-associated microorganisms. The current study focuses on the bacterial isolates, included those that originate from growth media supplemented with antibiotics.

The 16S rRNA gene of each isolated strain was sequenced using the primers 27F (5’-AGAGTTTGATCCTGGCTCAG-3’) and 1492R (5’-CGGTTACCTTGTTACGACTT-3’), and strains were identified by aligning this sequence on the NCBI database (Sayers et al., 2022). In the present study, the collection was split into two groups. All the strains from the Actinomycetes class belonged to the genera *Streptomyces*, *Nocardioides* and *Nocardia*, and will be referred to as the filamentous strains. All the other strains will be referred to as the non-filamentous strains.

Non-filamentous bacteria were routinely cultured by streaking a single colony onto a fresh plate of TSA agar plate and incubated at 28 °C for 1 to 5 days, depending on the strain. Cultures were maintained on the same plate for no longer than two weeks before being subcultured onto fresh TSA agar. The filamentous strains were routinely cultured by collecting spores from a fresh plate and streaking them onto a fresh Petri dish containing MV agar medium (10% Alnatura Gemüse Direktsaft supplemented with 1 g/L of CaCO_3_ and 15 g/L of agar). The inoculated plates were incubated for 7 days at 28 °C until spore formation before being stored in sterile distilled water at 4 °C for up to 1 year. For the cross-streak and round plate assays described below, centrifuged MV (cMV) was produced by centrifuging MV medium at 5’000 rpm for 10 minutes prior to adding the agar and CaCO_3_ to remove the largest tomato particles.

#### Origin and cultivation of the pathogen strains

*Pectobacterium carotovorum* Imp16/645-4 (Pcar), *Dickeya solani* (Dsol) CH16/120-3, *Alternaria solani* isolate 2713 (Asol) and *Pythium ultimum* isolate 212 (Pult) were provided by Agroscope (Nyon, Switzerland). *Rhizoctonia solani* strain 5900 was provided by Syngenta SA (Basel, Switzerland), and a *Phytophthora infestans* strain of genotype EU_44_A1 (Pinf) that was isolated from Reckenholtz, Switzerland in 2021.

The bacterial pathogen Dsol and Pcar were routinely cultured the same way as the non-filamentous strains from the collection. The fungal and oomycete pathogens were routinely grown by placing a 5 mm plug of mycelium on a fresh plate containing MV agar medium and incubating it at 28 °C for 7 to 10 days for Asol, 3 days for Rsol and Pult and 2 weeks at 21 °C for Pinf.

### Confrontation assays

#### Non-filamentous strains

##### Preparation of P. carotovorum and D. solani inoculates

Luria-Miller LB broth (Carl Roth, Karlsruhe, Germany) was inoculated with single colonies of Pcar or Dsol and grown under agitation (120 rpm) overnight at 28 °C. Bacterial cells were harvested by centrifugation at 4’500 x g for 5 minutes and washed twice in sterile distilled water. The clean bacteria suspension was mixed in molten MV agar medium at 50 °C to reach a final optical density at 600 nm (OD_600_) of 0.1. About 35 mL of inoculated MV agar was quickly spread as a thin layer at the surface of a square Petri dish (120 mm per 120 mm) containing MV agar medium and left to set for 1 h before use.

##### Preparation of ground P. ultimum, A. solani and R. solani mycelium inoculates

Ground mycelium was used for the confrontation assays against Pult, Asol and Rsol. For Pult, the mycelium from each plate was picked and transferred into four 2 mL Eppendorf tubes containing 1 mL of sterilized water and iron beads. For Asol and Rsol, a sterile cellophane sheet (Membrane backing model 583 gel dryer, Bio-Rad Laboratories, USA) was laid on a fresh plate of MV agar medium to facilitate mycelium recovery. The plates were inoculated with a 5 mm plug of fresh mycelium and were incubated at 18 °C for 2-3 weeks. The mycelium of each a plate was then carefully scrapped with a metal spatula. The mycelial material was distributed in 4 Eppendorf tubes (2 mL) for Asol and 6 tubes for Rsol. These numbers of tubes were determined empirically to provide homogeneous mycelial growth while avoiding overgrowth in the confrontation plates. One millilitre of sterile distilled water and sterile iron beads were added to each tube.

For the three pathogens, the harvested mycelium was ground by shaking the tubes at 25 beat per seconds for 1 minute and 30 seconds in a MM400 mixer mill (Retsch, Haan, Germany). The metal beads were removed, and the ground mycelium was added to molten MV agar and plated thinly on a square Petri dish containing MV agar as described above. To ensure homogeneous growth, each square plate was inoculated with the content of one Eppendorf tube of mycelium.

##### Preparation of P. infestans sporangia inoculate

Pinf was grown on fresh MV agar medium for 10 to 21 days at 20 °C until it produced a large quantity of sporangia. Mycelium was then scrapped off the agar and put into a 15 mL Falcon tube, to which sterile distilled water was added until reaching 2 mL. The tube was shaken vigorously to detach the sporangia from the hyphae. The solution was then filtered using a nylon mesh. The sporangia concentration was determined using a Thoma chamber and added to molten MV agar to reach a final concentration of 50’000 sporangia/mL. The inoculated molten MV agar medium was spread on a square Petri dish containing MV agar and left to set for 1 h before use.

### Bacterial growth rate measurement and preparation

The growth rate of each strain was measured from 96-well plates. Two hundred microliters of filtered liquid V8 medium (autoclaved 10 % V8 juice supplemented with 1 g of CaCO_3_, filtered using 22 μm sterile syringe filters) (Miller, 1955) was added to each well and inoculated with a single colony from a fresh culture. It was then incubated at 28 °C under agitation at 120 rpm for three days, and the OD_600_ of each well was measured twice per day using a Cytation5 cell imaging reader (Agilent, Santa Clara, USA). The strains were then grouped into batches of maximum 36 strains per 96-well plate (6 per row and per 6 column) according to the time they took to reach the stationary phase (up to 10 h, 10-20 h, 20-24 h, 24-35 h) and glycerol stocks were generated in this 96-well plate configuration. The glycerol stock was used to inoculate mother plates with 200 mL of fresh filtered MV medium. The mother plates were incubated at 28 °C under agitation at 120 rpm until they reached the stationary phase (10 h to 72 h depending on the growth speed plate). They were then used to inoculate a new 96-well plates (same growth medium and conditions) which served as inoculum for the confrontation assays between bacteria and pathogens, and stored up to one week at 4 °C.

### Confrontation plate inoculation and imaging

The inhibitory potential of the strains was assessed by confronting each batch of 36 strains against each pathogen (Figure 1a). Four microliters of each strain suspension were pipetted onto a fresh pathogen-inoculated square Petri dish. The suspensions were pipetted per columns (of each six row), and an adjustable-spacing multichannel pipette was used to maximize the distance between the resulting colonies on the destination square Petri dishes.

**Figure 1:**
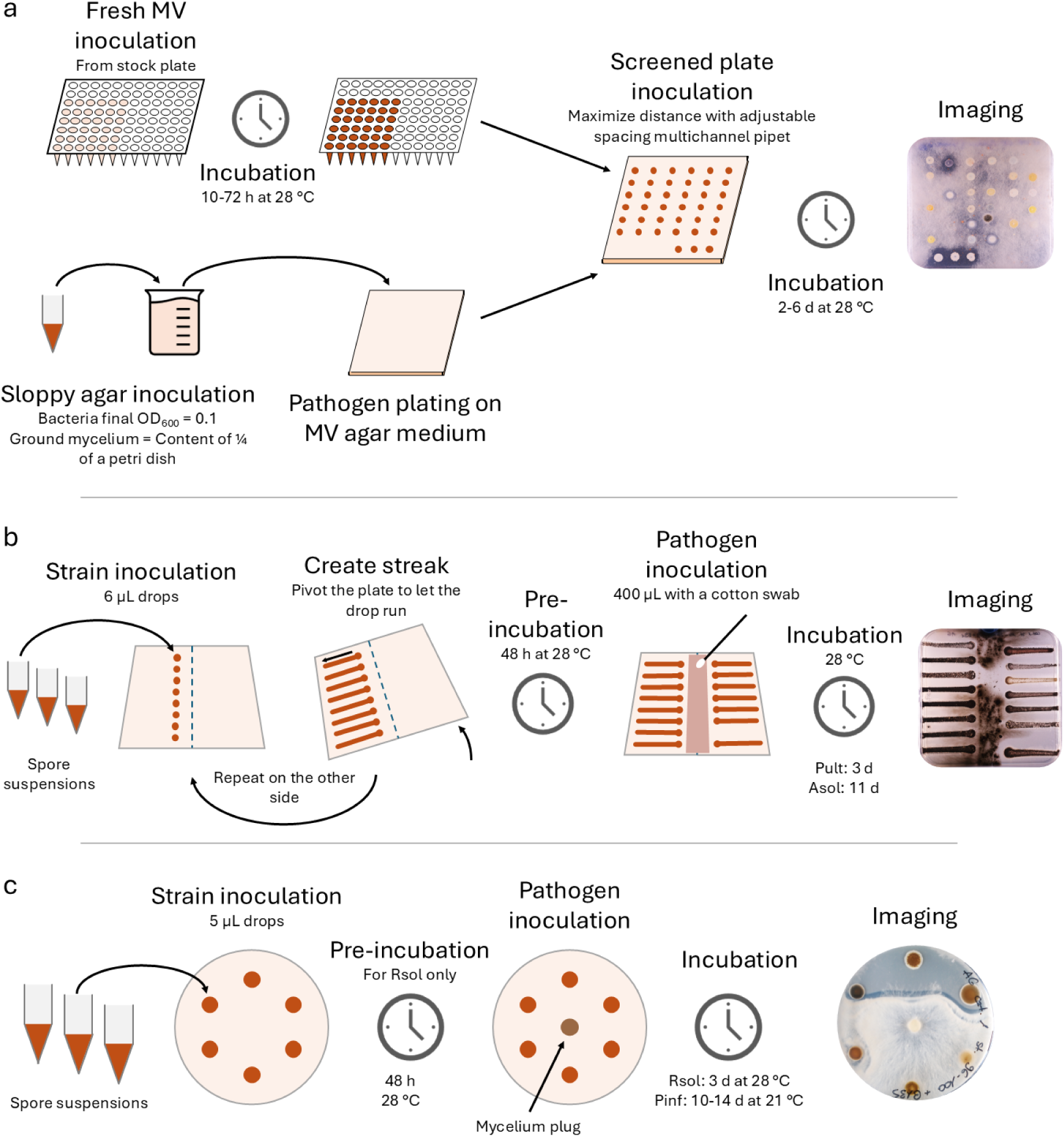
**(a)** Screening of the activity of the non-filamentous strains against the pathogens *Dickeya solani* (Dsol), *Pectobacterium carotovorum* (Pcar), *Alternaria solani* (Asol), *Rhizoctonia solani* (Rsol), *Phytophthora infestans* (Pinf) and *Pythium ultimum* (Pult). Fresh MV medium in 96-well plates was inoculated with up to 36 strains of similar growth rate and incubated until maximal growth was reached. Cell suspensions or ground mycelium were mixed in soft agar at 50 °C and poured as a thin overlay onto square agar plates containing MV. Four-microlitre droplets of each bacterial culture were then spotted onto the overlay using an adjustable-spacing multichannel pipette to maximize the distance between colonies. Pathogen-and collection strain-inoculated plates were incubated for 2–6 days, depending on pathogen growth rate, before imaging. **(b)** Screening of the activity of the filamentous strains against Dsol, Pcar, Asol and Pult. Spore suspensions were stored in separate tubes to prevent cross-contamination. Six-microlitre droplets were streaked onto fresh centrifuged MV (cMV) agar by tilting the plate immediately after inoculation. After 48 h pre-incubation, the pathogen inoculum was applied as a central band between the bacterial streaks. Pathogen-and collection strain-inoculated plates were then incubated for 24 h (Dsol, Pcar), 3 days (Pult) or 11 days (Asol) before imaging. **(c)** Screening of the activity of the filamentous strains against Rsol and Pinf. Five-microlitre droplets of six filamentous strains were spotted onto fresh cMV agar in a circular arrangement. A fresh mycelial plug was placed at the centre of the plate either immediately after bacterial inoculation (Pinf) or following a 48 h bacterial pre-incubation (Rsol). Pathogen-and collection strain-inoculated plates were then incubated for 3 days (Rsol) or 10-14 days (Pinf) before imaging.

The strain-inoculated Petri dishes were allowed to dry until the droplets were completely absorbed by the agar and subsequently incubated according to the pathogen used. The plates inoculated with Dda and Pcar were incubated for 1 day at 28 °C; the plates inoculated with Asol and Rsol were incubated for 3 days at 20 °C; the plates inoculated with Pinf were incubated for 4 to 6 days at 20 °C; and the plates inoculated with Pult were incubated for 2 days at 20 °C. Following incubation, both sides of each plate were photographed to visualize the inhibition areas. For each assay, two control plates were incubated alongside the confrontation plates: one containing only the pathogen and one containing only the tested strains.

#### Filamentous strains

##### Preparation of the pathogen inocula

The bacterial pathogens Dsol and Pcar were prepared as described for the non-filamentous strains until the cell harvest. After cleaning, the pellet was diluted in distilled water to reach an OD_600_ of 1. The ground mycelium inocula of Asol and Pult were prepared as described for the non-filamentous strains. For Pinf and Rsol, five millimetres plugs of mycelium were punched from mature plates that were less than 2-weeks old.

##### Isolate preparation

For each strain, spores from a single colony were picked with a loop and incubated on a new plate containing MV agar medium for 7 days at 28 °C. A volume of 5 mL of sterile distilled water was poured onto the plate and the aerial mycelium was scratched with a sterile loop. The mycelium-water solution was poured into a Falcon tube and vigorously shaken to detach the spores from the mycelium. The solution was then filtered through a sterile filter made of a glass wool-filled 10 mL pipette tip into a new Falcon tube. The resulting spore suspensions were adjusted to OD_600_=1 and stored at 4 °C.

##### *Cross-streak on* D. solani, P. carotovorum, A. solani*, and* P. ultimum

Filamentous strains were screened with two methods to fit the pathogens growth specificity. The first method was a modified cross-streak (Figure 1b) as described in (Gillon et al., 2023). A square Petri dish with fresh centrifuged MV agar was inoculated with fifteen filamentous strains. A vertical line was drawn at the centre of the plate, and 6 µL drops of spore suspension from 8 strains were placed 1.5 cm on the left of the line. The drops were separated from each other by 1.2 cm. Immediately after pipetting, the plate was rotated to the left for the drops to migrate towards the edge of the plate, leaving a streak. The same process was repeated on the right side of the line with 7 strains, including the negative and positive controls (i.e. strains of known activity vs. inactivity against the respective pathogens), which were placed at a 2.4 cm distance from each other. The plates were then incubated at 28 °C for 48 hours before pathogen inoculation.

For the bacterial pathogens, a cotton swab was used to apply 400 µL of the cell suspension as a 2 cm-thick band that followed the centre line, while staying 5 mm away from the filamentous bacteria streaks. The plates were incubated at 28 °C for 24 hours before imaging.

The same process was used for the fungal pathogen Asol and oomycete pathogen Pult but using 400 µL of the ground mycelium inoculum instead. The plates were incubated at 28 °C for 11 days for Asol and 3 days for Pult before imaging.

##### Round plate assay for P. infestans and R. solani

A round plate assay was used to assess the inhibitory activity of the filamentous strains on Pinf and Rsol. A 9-cm Petri dish with cMV medium was inoculated with 5 µL drops of 6 filamentous bacteria placed at 3.5 cm distance to each other and to the centre of the plate. For Pinf, the fresh mycelium plug was immediately placed at the centre of the plate. Plates were then incubated at 21 °C for 10-14 days before imaging. For Rsol, the plate with the isolates was preincubated at 28 °C for 48 hours before adding the fresh mycelium plug and subsequently incubated at 28 °C for 3 days before imaging.

### Scoring of activity and data analysis

#### Inhibitory activity scoring

The inhibitory potential of each strain against a single pathogen was assessed by manual scoring. A score of 0 indicated no inhibition. A score of 1 indicated a weak inhibition, where pathogen growth stopped at the edge of the bacterial colony. A score of 2 indicated a moderate inhibition, where pathogen growth stopped between 1 and 5 mm from the edge of the colony. A score of 3 indicated a strong inhibition, where pathogen growth stopped at more than 5 mm from the edge of the colony, with the inhibition zone sometimes extending to neighbouring colonies. A score of 4 indicated an outstanding inhibition, where pathogen growth was prevented over a sufficiently large area to entirely encompass neighbouring colonies (Figure S1).

An observation was considered invalid if a score could not be assigned due to a contamination, interference from an adjacent outstanding inhibition area, or if the tested strain failed to grow in both the confrontation plate and the strain control plate. At least two replicates were performed for each strain-pathogen combination. From this point, each combination of a strain and a pathogen will be referred to as a “confrontation”. Additional replicates were performed when only one replicate was valid, or when the score difference between the first two replicates of a confrontation exceeded one, as such cases were considered unreliable.

#### Data cleaning and confrontation categorization

The data was analysed using R v4.5.2 (R Core Team, 2021) running from RStudio build 218 (Posit team, 2025).

The data was filtered to remove strains and confrontations with repeated invalid output. Strains for which fewer than 10 valid observations across all pathogens were obtained were excluded, as they were considered unsuited for the current method. Confrontations with only one valid observation, as well as the confrontations with 2 valid observations showing a score difference greater than one, were also removed because they were considered unreliable.

Activity categories were assigned based on median confrontation scores: inactive (<1), weakly active (1 to <2), moderately active (2 to <3), highly active (3 to <4), and outstanding (=4). While this classification allowed direct comparison of all the strains, the filamentous strains will be analysed separately from the remainder of the collection because they were tested with distinct protocols.

#### Proportion of strain activity per isolation characteristics and per pathogen

The influence of isolation characteristics on activity was investigated. The strains were grouped according to their cultivar of origin, their compartment of origin or their isolation medium. Each strain was assigned the highest activity category it achieved against any pathogen, and the distribution of these highest activity categories was compared among the isolation characteristics groups.

#### Annotated phylogenetic tree construction

Two phylogenetic trees were generated, one for the filamentous strains one for the non-filamentous strains, based on their partial 16S rRNA sequence. Sequences were aligned with MAFFT v7.505 (Katoh and Standley, n.d.), before being trimmed with TrimAl v1.5 (Capella-Gutiérrez et al., 2009) at 95% coverage. This procedure created aligned sequences of 974bp for the non-filamentous strains and 768bp for the filamentous strains. Phylogenetic trees were computed using FastTree v2.1.11 (Price et al., 2010).

To improve readability, the phylogenetic trees were rescaled with a delta model using the ape v5.8-1 (Paradis and Schliep, 2019) package. The trees were visualized and annotated using the ggtree v4.0.4 (Xu et al., 2022) and ggtreeExtra v1.20.1 (Xu et al., 2021) packages.

#### Activity of the main genera per pathogen

The genera that were represented by at least 10 strains in the collection, including at least 15% of active strains against one pathogen were selected for further analysis. For each selected genus, the proportion of active strains relative to the total number of strains within that genus was calculated for each pathogen. For each selected pathogen, the proportion of active strains per genus was calculated relative to the total number of active strains for the respective pathogen.

## Results

### Development of high-and medium-throughput screening methods

A single method allowed the fast screening of the inhibitory activity of 363 non-filamentous strains, belonging to 57 different genera, against six different pathogens (Figure 1a). From the original 391 non-filamentous strains, 28 were removed because they grew poorly with the described method. From the remaining strains, 13 combinations of a strain and a pathogen (referred to as “confrontations”) were removed because the pathogen inhibited the growth of the strain. In total, the inhibitory activity was successfully assessed in 2’163 confrontations out of 2’346 possible, which represents a 92% success rate.

Filamentous strains had to be tested using two different protocols due to their slower growth speed and higher inhibitory potential (Figure 1b-c). These lower-throughput methods and the greater homogeneity of these strains returned more reliable results, with a success rate of 97%, leaving only 3 strains belonging to the *Streptomyces* genus, 2 *Micromonospora*, one *Promicromonospora*, and one *Nocardia* untested. While the scoring scale used was the same for filamentous and non-filamentous strains, these two groups will be discussed separately as different methods were used.

The screening of such a high quantity and diversity of strains that originated from the same plants allowed further analysis on the parameters influencing inhibitory potential.

### Isolation characteristics had a marginal influence on inhibitory activity

The collection comprised 600 strains, isolated in approximately equal proportion between the two cultivars Innovator and Bintje. The highest number of strains was isolated from the rhizosphere soil (263) and the root (231), much less strains originating from the phyllosphere (106). A similar proportion of strains were isolated from 10% TSA, ARE and ActIA media (205, 176 and 204 respectively), and PDA had a far lower number of isolates (15). Taxonomically, the isolated strains belonged to 62 genera from four phyla: 362 Actinobacteria, 83 Firmicutes, 137 Proteobacteria and 18 Bacteroidota. About a third of the isolated strains (208) were filamentous actinomycetes.

The influence of the compartment, cultivar and medium of isolation on the inhibitory potential of the isolated strains was evaluated (Figure 2). The filamentous strains from the collection had a far stronger inhibition potential than the non-filamentous ones, with 74% of the strains that were active against at least one pathogen, compared with 26% for the non-filamentous ones.

**Figure 2:**
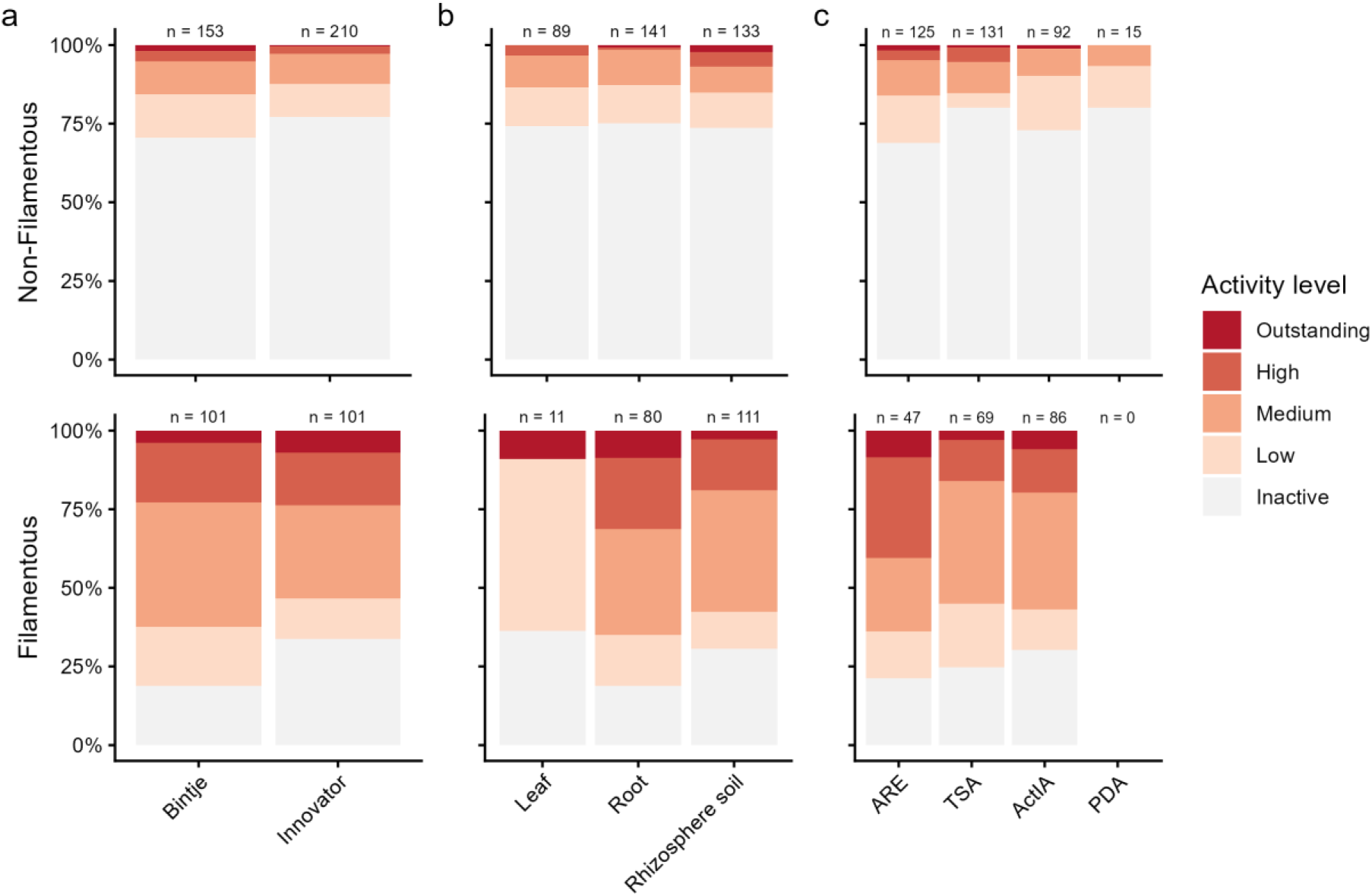
Distribution of the maximum antagonistic activity of bacterial isolates according to cultivar of origin (a), plant compartment of origin (b), growth medium used for isolation (c), and strain type (non-filamentous vs. filamentous). Stacked bar plots show the proportion of strains assigned to each activity level (inactive, low, moderate, high, and outstanding), with each bar representing 100% of the strains within a given category. For each strain, the activity level corresponds to the highest level of antagonism observed against any of the tested pathogens. For example, a strain exhibiting moderate activity against a single pathogen and no activity against the others was classified as moderately active.

No striking influence of the cultivar, compartment or medium of isolation on the distribution of antagonistic activity were observed, but some interesting trends still emerged:

Comparing both cultivars, we observed a slightly higher frequency of active strains coming from the susceptible cultivar Bintje than from the resistant cultivar Innovator among both non-filamentous and filamentous strains (Figure 2a).

The compartment of isolation did not influence the proportion of active non-filamentous strains, with almost identical proportions in the leaf-, root-and rhizosphere soil-isolated strains. However, the highest activity levels were observed in the rhizosphere soil, from where most of the strains showing outstanding activity were isolated (2.3%). This compartment also displayed the highest proportion of highly active strains with 4.5% in the rhizosphere soil, compared to 3.4% and 0.7% in the leaf and root respectively (Figure 2b). Different tendencies were observed in the filamentous strains, where most of the strains isolated from roots and rhizosphere soil displayed an activity of medium level or higher (65% and 57%, respectively), while leaf isolates showed either outstanding (9%) or low (54%) activity.

The impact of the growth media on inhibitory activity was similar between filamentous and non-filamentous strains. The ARE growth medium provided the highest numbers of active strains in both filamentous and non-filamentous strains (78% and 31%, respectively) and included most of the strains with an outstanding activity. In comparison, the TSA and ActIA media yielded a lower but still substantial proportion of active strains (27% and 20% for the non-filamentous and 75% and 70% for the filamentous, respectively), with lower activity levels. In contrast, PDA did not seem to foster the isolation of particularly active strains in non-filamentous bacteria, while no filamentous strains could be retrieved from it.

Despite these tendencies, the influence of the different isolation characteristics on antagonistic activity remained limited overall. While analysing each pathogen separately did not reveal additional patterns (Figure S2), it showed that the pathogens had different resistance levels to growth inhibition by potato-associated bacteria, and that the high activity levels observed among the filamentous strains was essentially driven by the hypersensitivity of a single pathogen to bacterial inhibition.

### Pathogens from different kingdoms differed in their sensitivity to bacterial antagonism

The proportion of active strains per pathogen was analysed. The bacterial pathogens *D. solani* (Dsol) and *P. carotovorum* (Pcar) were remarkably resistant against the inhibitory activity of the tested bacterial strains. Only 3% of the filamentous strains and 0.6% of the non-filamentous strains displayed inhibitory activity against Dsol (Figure 3). Similar results were observed for Pcar with 3.5% of active filamentous strains and 0.8% of the non-filamentous strains (Figure 3). While the activity of non-filamentous strains against these pathogens was medium at best, a few filamentous strains still displayed a high inhibition activity.

**Figure 3:**
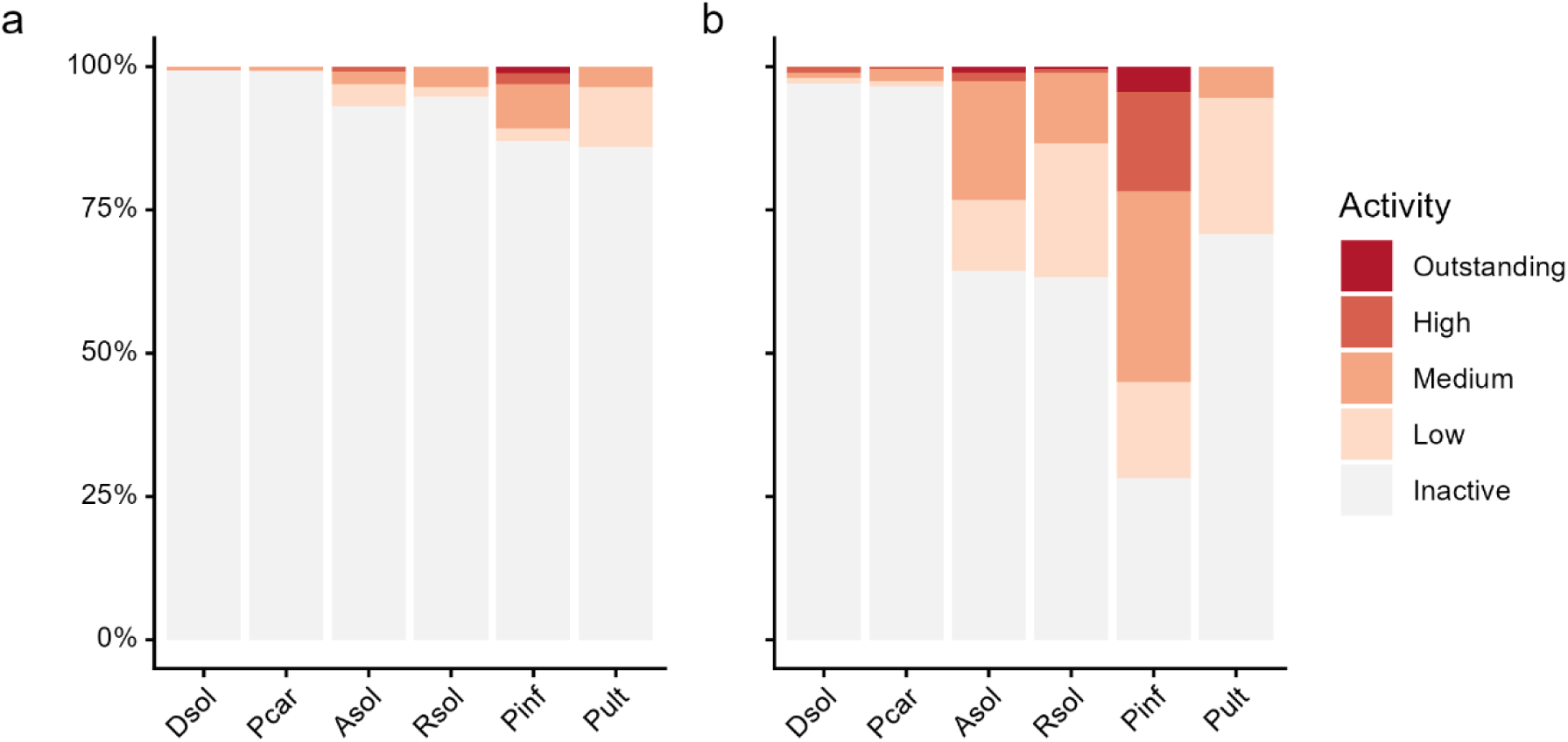
Distribution of antagonistic activity levels of non-filamentous **(a)** and filamentous **(b)** bacterial isolates against each pathogen. Stacked bar plots show the proportion of strains exhibiting each activity level (inactive, low, moderate, high, and outstanding) for each of the six tested pathogens. Each bar represents 100% of the strains tested against the corresponding pathogen.

In comparison, the fungal pathogens were more sensitive to bacterial antagonism. Although *A. solani* (Asol) and *R. solani* (Rsol) had similar inhibition frequency (∼6% for non-filamentous strains, ∼36% for filamentous strains), Asol displayed a higher sensitivity to filamentous strains than Rsol, with a higher proportion of strains showing medium, high and outstanding levels of activity (23% vs 13%) (Figure 3). Both fungal pathogens had a similar sensitivity to non-filamentous strains, with a lower proportion of moderately active strains against Asol than against Rsol (2.2% and 3.6%), compensated by the presence of a few highly active strains against Asol (0.8%) (Figure 3a).

The oomycete pathogens had contrasting sensitivity to bacterial antagonism. *P. infestans* (Pinf) displayed by far the highest sensitivity to the strains of all pathogens. This sensitivity was the most striking when confronted with the filamentous strains, 72% of which had an antagonistic activity against this pathogen. On top of this high frequency, the inhibition observed was also intense, with 33% of medium activity, 17% of high activity, and an exceptionally high 4.5% of outstanding activity (Figure 3b). The same sensitivity was observed when Pinf was confronted to the non-filamentous strains, but to a lower extent, with 13% of antagonistic strains, among which a high proportion of medium (7.7%), high (1.9%) and outstanding (1.1%) intensity inhibitions (Figure 3a). The second oomycete, *P. ultimum* (Pult), showed comparable sensitivity towards non-filamentous strains (14%), but with far lower intensity (10% of low and 3.6% of medium activity). However, among the filamentous strains, the proportion of inhibition was lower than that observed against Pinf (72%) and against the fungal pathogens (29%) (Figure 3b).

While these analyses describe overall pathogen sensitivity at the collection level, they do not indicate the phylogenetic identity of active strains nor whether inhibition of different pathogens was mediated by similar or different sets of strains, which was the next aspect we investigated.

### Most active strains clustered within a few orders

The inhibitory activity of each isolate against each pathogen was mapped onto phylogenetic trees to assess whether antagonistic phenotypes clustered with phylogenetic relationships. The identities of the active strains within these clusters are provided in Figure S3.

Among non-filamentous isolates, most active strains tended to group into distinct phylogenetic clusters (Figure 4a). The strongest inhibitory activity was observed within a cluster belonging to the Bacillales order (bright blue colour). This cluster contained multiple strains capable of inhibiting Asol, Rsol, and Pinf, as well as two of the only three strains able to inhibit bacterial pathogens. Notably, this was also the only cluster containing strains capable of inhibiting both Dsol and Rsol. In addition, it was the only cluster harbouring non-filamentous generalist inhibitory strains. Surprisingly, however, Pult was not affected by strains belonging to this cluster.

**Figure 4:**
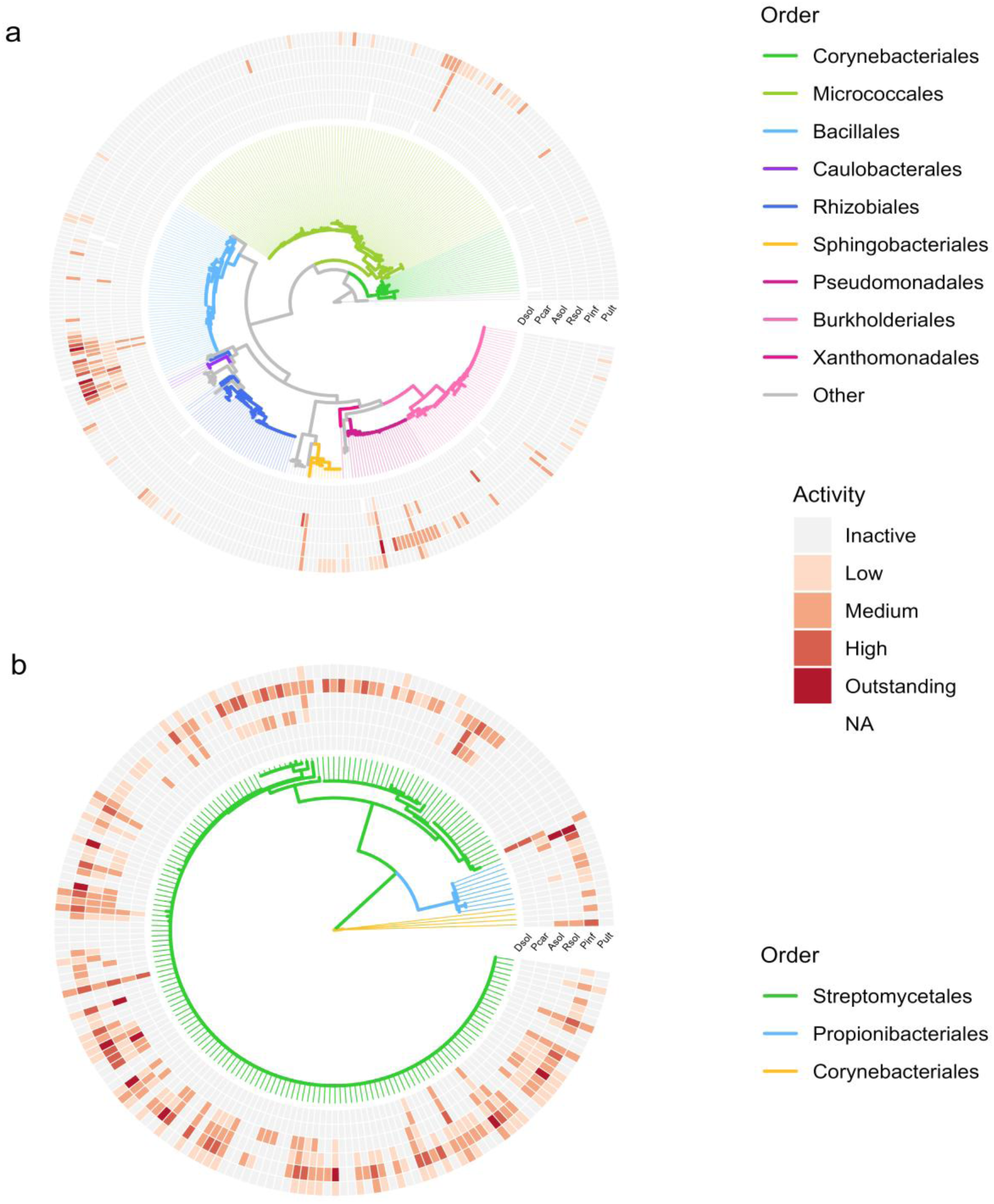
Phylogenetic trees showing the antagonistic activity of each bacterial isolate against the six tested pathogens for non-filamentous **(a)** and filamentous **(b)** strains. Branch colours indicate the taxonomic affiliation of each isolate, corresponding to the order of the strains. The concentric heatmap surrounding each tree represents the antagonistic activity level of each strain against each pathogen. White colour represents missing data. From the innermost to the outermost ring, activity is shown against *Dickeya solani* (Dsol), *Pectobacterium carotovorum* (Pcar), *Alternaria solani* (Asol), *Rhizoctonia solani* (Rsol), *Phytophthora infestans* (Pinf), and *Pythium ultimum* (Pult).

In contrast to the other pathogens, Pult was almost exclusively inhibited by specialist inhibitory strains. These strains formed a large cluster within the Micrococcales (olive green), as well as smaller clusters within the Sphingobacteriales (yellow), Rhizobiales (dark blue), and Pseudomonadales (deep pink) and Burkholderiales (pink) (Figure 4a). An additional cluster within the Pseudomonadales harboured strains with high Pinf-inhibitory activity, with two strains causing general inhibition of all non-bacterial pathogens.

The phylogenetic tree of filamentous isolates consisted of strains belonging to three orders: Streptomycetales (green colour, all from the *Streptomyces* genus), Propionibacteriales (blue colour, all from the *Nocardioides g*enus), and Corynebacteriales (yellow colour, all from the *Nocardia* genus) (Figure 4b). Most isolates from this group belonged to the Streptomycetales order, a single branch of which occupied approximately two thirds of the tree (from approximately 3 o’clock to 11 o’clock), reflecting strains sharing identical partial 16S rRNA gene sequences. However, this apparent phylogenetic similarity did not result from strain redundancy, as these isolates displayed distinct colony morphologies and inhibitory activity profiles.

This large Streptomycetales branch contained numerous generalist inhibitory strains capable of inhibiting all fungal and oomycete pathogens tested. Five of these strains were further able to inhibit all tested pathogens, including bacterial pathogens, a characteristic not observed among non-filamentous isolates. However, the branch also contained completely inactive strains and only a small number of specialist inhibitory strains.

The remaining Streptomycetales isolates exhibited more distinct activity-associated clustering. From approximately the 11 o’clock to the 2 o’clock position on the phylogenetic tree, a cluster of strains capable of inhibiting both Asol and Pinf was followed by a cluster of specialist Pinf inhibitors, and subsequently by two small clusters of generalist inhibitory strains separated by a group of inactive isolates. In contrast, the few isolates belonging to the Propionibacteriales and Corynebacteriales orders displayed overall low inhibitory activity besides Pinf antagonism.

### A closer look at highly active bacterial orders

As a final step, the bacterial orders that contained the highest proportion of inhibiting strains for each pathogen were analysed in more details. The proportion of strains displaying inhibitory activity relative to the total number of isolates within a given order will be referred to as the order activity prevalence (shown as percentage of active strains in each order for each pathogen in Figure 5a), and the relative representation of orders among all strains active against each pathogen will be referred to as the order composition of active strains (shown as order distribution of active strains for each pathogen in Figure 5b). For more details, the distribution of the activity levels of the predominant orders of the bacterial collection against each pathogen can be found in Figure S4.

**Figure 5:**
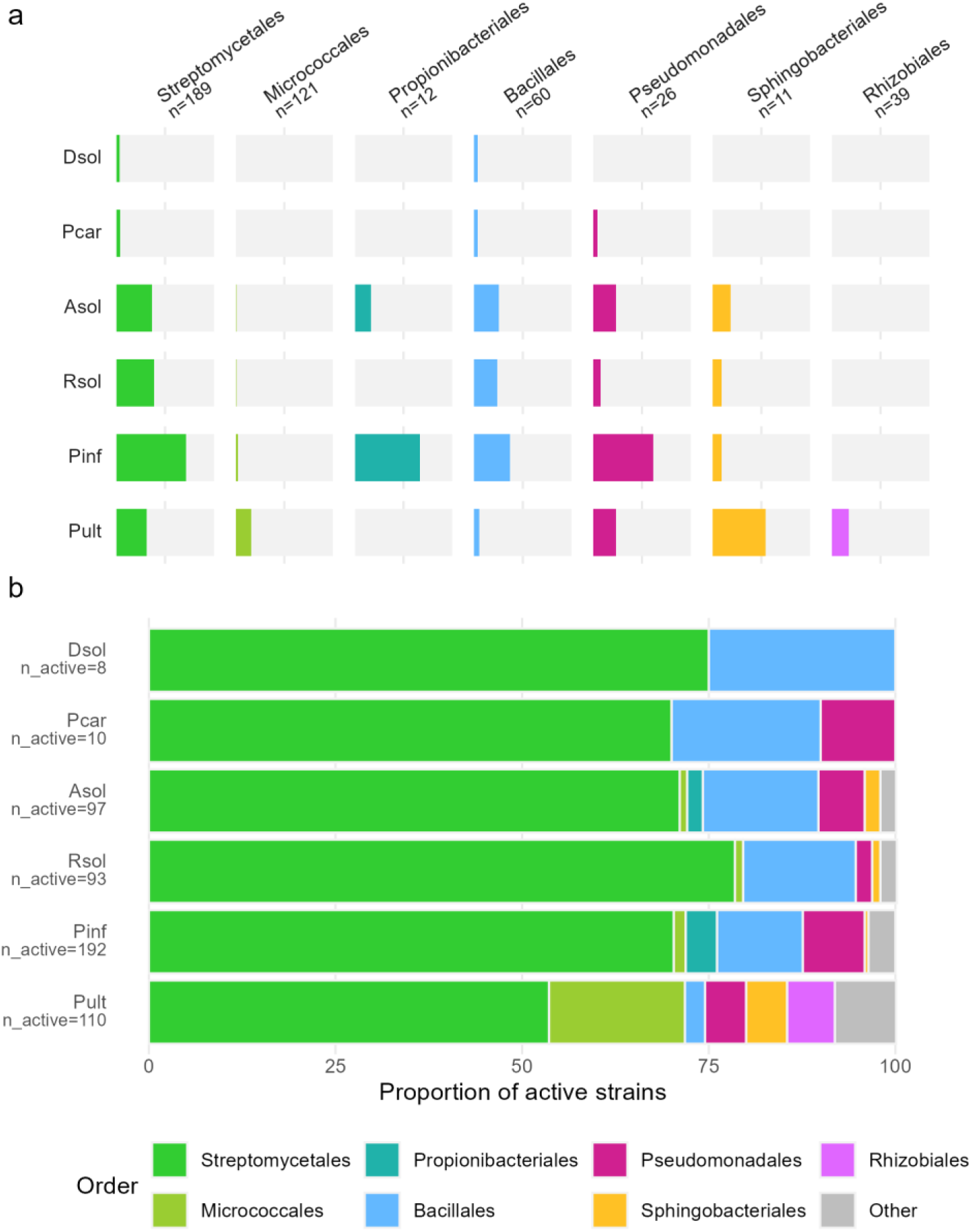
Distribution of active strains among the predominant bacterial orders in the collection. Only orders containing genera represented by at least 10 strains and exhibiting at least 15% active strains were included. Active strains were defined as strains displaying at least low antagonistic activity against the corresponding pathogen. **(a)** Proportion of active strains within each bacterial order for each pathogen, represented as horizontal bar plots faceted by bacterial order and pathogen. **(b)** Taxonomic composition of active strains for each pathogen. Each stacked bar represents the complete set of active strains against a given pathogen, with segments indicating the relative contribution of each bacterial order to the active strain pool.

The bacterial pathogens Dsol and Pcar had a very low number of antagonistic strains (8 and 10, respectively) (Figure 5b) and a similar order activity prevalence that included a small fraction of the Streptomycetales and Bacillales (∼3.5%) (Figure 5a). In addition, a small fraction of the Pseudomonadales were active against Pcar but not against Dsol.

The phylogenetic composition of active strains was similar between the fungal pathogens, for which the number of active strains, with 97 for Asol and 93 for Rsol, was more than 10-fold higher than that observed for the bacterial pathogens (Figure 5b). The order activity prevalence of the fungal pathogens was similar for the Streptomycetales and Bacillales (about 37% and 25% for both), but Asol was more sensitive than Rsol to the Pseudomonadales (23% against 8%, respectively) and to additional orders such as the Sphingobacteriales (about 18% against 9%) and the Propionibacteriales (16.7% against Asol only). A small fraction of the Micrococcales were also antagonistic to both fungal pathogens (<1%) (Figure 5a).

The oomycete pathogens each had a unique inhibition pattern. Pinf was inhibited by the highest number of strains (192) (Figure 5b). The same orders that were active against Asol were also active against Pinf, but with a far higher activity prevalence among the Streptomycetales, Propionibacteriales, Pseudomonadales and Bacillales (71%, 66%, 61% and 37% respectively), and a lower activity prevalence among the Sphingobacteriales (9%) (Figure 5a). The composition of strains active against Pinf was composed of similar proportions of Streptomycetales and Micrococcales as that active against the fungal pathogens (70% and 1.6%), of a higher proportion of Pseudomonadales and Propionibacteriales (8% and 4%), and of a lesser proportion of Sphingobacteriales (<1%) (Figure 5b). Lastly, the inhibition pattern of Pult markedly differed from all the other pathogens. It was the only pathogen inhibited by Rhizobiales, with an activity prevalence of 17%. In addition, the order activity prevalences of the Micrococcales and Sphingobacteriales were higher than in any other pathogen (16% and 54%), but those of the Streptomycetales and Bacillales were lower than for the other oomycete and fungal pathogens (31% and 5%) (Figure 5a). In the end, these differences strongly affected the active strain composition of Pult, with the lowest proportion of Streptomycetales and Bacillales (54% and 2.7%, respectively), and by far the highest proportions of Micrococcales, Rhizobiales and Sphingobacteriales (18%, 6.4% and 5.5%, respectively), compared to all other pathogens (Figure 5b).

## Discussion

In this study, the antagonistic activity of 600 strains isolated from potato plants was screened against 6 potato pathogens: the bacterial pathogens *Dickeya solani* (Dsol) and *Pectobacterium carotovorum* (Pcar), the fungal pathogens *Alternaria solani* (Asol) and *Rhizoctonia solani* (Rsol) and the oomycetes *Phytophthora infestans* (Pinf) and *Pythium ultimum* (Pult). A single method was used for all the non-filamentous strains, and two methods were used for the filamentous strains.

Most screenings involving direct pathogen confrontations are tedious and time-consuming, with the workload scaling with the product of the number of strains screened by the number of pathogens tested (Berg et al., 2002; Caulier et al., 2018; Guyer et al., 2015). Although methods that reduced the workload of this process have been developed, they either did not substantially reduce scalability (Upreti and Thomas, 2015), were not adapted for fungal pathogens (Anzalone et al., 2021), or relied on supernatant filtration (Munakata et al., 2022), thereby preventing direct contact between the tested strain and the pathogen.

One of the major aims of this study was to develop a method capable of screening a broad range of strains against a broad range of pathogens, while remaining easily scalable and accessible to most microbiology laboratories. The method presented here for the non-filamentous strains meets all these criteria through two key features. Firstly, organizing the strain collection into groups of 36 strains of similar growth rates ensured consistent results by minimizing the number of invalid confrontations due to fast-growing or spreading strains. Storing these groups as glycerol stocks adapted for long-term storage allowed them to be readily replicated into fresh plates. This reduced the risk of contamination associated with repeated reinoculations into new plates, while saving time when adding new pathogens or performing additional replicates. Secondly, incorporating the pathogen inoculum directly into molten agar at low temperature proved adaptable to all six pathogens tested, with minimal modification. While this approach was naturally suited to bacterial pathogens or pathogens that sporulate readily *in vitro* such as Pinf, ground mycelium had to be used for the other pathogens. The mycelium of *P. ultimum* (Pult) could be readily harvested from the growth medium, whereas for fungal pathogens whose mycelium strongly adheres to agar, a sterilized cellophane sheet placed between the growth medium and the mycelial plug enabled efficient harvesting.

Although effective, this method was not suitable for the filamentous strains of the collection. Those were stored as spore suspensions, which were prone to airborne cross-contamination, and therefore could not be organized into multi-well strips or plates. In addition, their slower growth rates prevented them from being inoculated onto the pathogen-inoculated agar overlay. Consequently, the cross-streak and conventional round plate methods were used, testing 13 and 5 strains per plate, respectively with their controls.

Although these methods enabled a large amount of data to be generated efficiently, they required particularly thorough assay validation, as their higher throughput also makes them more susceptible to contaminations between strains from neighboring wells compared to approaches that scale less efficiently. Moreover, one limitation of such screenings is that strains inoculated on the same plate likely have an influence on the metabolites produced by neighboring strains. Such influence could be reduced when randomizing the strain position between different biological replicates. Ultimately, the large dataset generated by these methods allowed us to investigate the influence of isolation parameters and phylogeny on antagonistic activity within our collection.

### Impact of isolation characteristics on antagonistic activity

The effect of the cultivation medium used for isolation showed a similar trend in filamentous and non-filamentous strains, with marginally higher proportions of active strains isolated from the ARE medium, which is the least rich medium we used (Figure 2). This result contrasts with the observations of a larger screening on 6’732 strains isolated from the rhizosphere of strawberry, rapeseed and from bulk soil, where the enrichment of rhizosphere suspensions with high molecular weight nutrients prior to plating on R2A growth medium systematically increased the proportion of isolated strains antagonistic to *Verticilium dahliae* (Berg et al., 2006). While the influence of the isolation medium on the proportion of active strains retrieved remains largely unresolved, as it is rarely documented in the literature, our observations indicate that a medium mimicking the composition of the rhizosphere such as the ARE medium, whose carbon sources are limited to few sugars, organic and amino acids rather than containing complex peptone and yeast extracts (Schmidt et al., 2016a), might be a good choice to isolate strains of relevance for the biological control of pathogens.

Plant genotype mildly affected the proportion of antagonistic strains, with a slightly higher proportion isolated from the susceptible cultivar Bintje for both filamentous and non-filamentous strains (Figure 2). This observation contrasts with findings from our previous study on the same collection, where a higher proportion of strains capable of inhibiting Pinf spore germination was isolated from the more resistant Innovator cultivar (Pichon et al., 2026). Other studies reporting the effect of the plant genotype on the proportion of antagonistic strains harboured are scarce. While a strong influence of plant species was observed following the screening of 5,854 strains against *V. dahliae* (Berg et al., 2002), contradictory observations were made at the cultivar level. In smaller scale screenings, strikingly similar proportions of *Bacillus* strains antagonistic to *V. dahliae* were found on leaf-isolated from 10 cultivated and 9 wild olive trees across Southern Europe (Müller et al., 2015), while a higher proportion of strains antagonistic to *Ralstonia solanacearum* was isolated from roots of tomato cultivars resistant to this pathogen (Upreti and Thomas, 2015). Overall, the question remains open of whether disease-resistant cultivars mainly rely on their plant genome-encoded determinants of resistance, with lesser dependence on associated microbiota, or whether part of their resistance stems from microbiota enriched in bacteria with high protective efficacy.

Similarly to the effect of the genotype, the influence of the isolation compartment on the proportion of antagonistic strains has been little studied so far and has yielded contradictory results: a screening of 424 strains isolated from tomato roots did not reveal any difference in proportions of strains antagonistic to several bacterial and fungal pathogens between the rhizosphere, rhizoplane, and root endosphere (Anzalone et al., 2021). However, a larger screening of 2’684 strains isolated from both surface and interior tissues of potato roots and leaves against *R. solani* and *V. dahliae* revealed higher proportions of active strains coming from the root endosphere and leaf surface than from the rhizosphere soil and leaf endosphere (Berg et al., 2005). In the current study, we found similar proportions of active strains between the compartments, but higher levels of activity among the non-filamentous strains isolated from the rhizosphere soil, and among the filamentous strains isolated from the roots (Figure 2). Interestingly, we found no association between the isolation compartment of the bacterial strains and the plant compartment targeted by the pathogen: root-and leaf-isolated strains were similarly capable of antagonizing pathogens originating from either compartment (Figure S2b). This suggests that the whole plant may be a valid source of potential mBCAs as antagonistic activity was not restricted to interactions between microorganisms originating from the same plant compartment.

The specificity of these antagonistic interactions across the pathogen panel was then examined to determine whether different pathogens exhibited distinct sensitivities to bacterial antagonism and whether specific taxa showed preferential activity against particular pathogens.

### Bacterial and fungal pathogens were inhibited almost exclusively by isolates with broad-spectrum activity

Most published screenings involving several pathogens show that their sensitivity to bacterial inhibition varies widely (Caulier et al., 2018; Guyer et al., 2015; Liu et al., 2017). The current study confirms this trend and adds that their sensitivity is linked to their phylogenetic proximity. The bacterial pathogens were the most resistant, with relatively weak antagonism observed for only about 1.5% of the collection, while the fungal pathogens were antagonised by about 16%, Pult by 19% and Pinf by 32% with far higher activity levels (Figure 3).

Most pathogen-inhibiting strains were grouped into clusters of phylogenetically closely related taxa (Figure 4). Among them were clusters of strains with broad-spectrum activity, so-called “generalist inhibitors”, that inhibited the growth of several pathogens, and clusters of more “specialist inhibitors” strains, that antagonize only one or two of them.

The generalist inhibitors clusters belonged to the Bacillales and Streptomycetales orders (Figure 4). More specifically, all the strains from the Bacillales cluster belonged to the *Bacillus* genus, which is almost systematically the most active genus in other screenings (Anzalone et al., 2021; Berg et al., 2005; Caulier et al., 2018; Müller et al., 2015). For this reason, the potential of *Bacillus* strains as biocontrol agent is widely studied (Dimkić et al., 2022), and 9 strains were approved in the European Union (EU) as biocontrol agents against diseases, with an additional 4 being assessed (“EU Pesticides Database - Active substances,” 2026).

The strains of the Streptomycetales generalist inhibitor cluster all belonged to the *Streptomyces* genus, which was the taxon with the highest number of inhibitory strains consistently across all pathogens (Figure 5b). This taxon is extensively studied for the high quantity and diversity of antibiotics and antifungal compounds they produce and for their potential as biocontrol agents (Alam et al., 2022). Two strains were approved as mBCA in the EU, and one is pending (“EU Pesticides Database - Active substances,” 2026).

Strains belonging to these Bacillales and Streptomycetales generalist inhibitor clusters contained almost all the strains capable of inhibiting the bacterial and fungal pathogens. Notably, the bacterial pathogens were only inhibited by generalist inhibitor strains, while a small number of specialists inhibitors were found for each fungal pathogen (Figure 4), mainly among the Burkholderiales, Pseudomonadales and Streptomycetales.

### The oomycete pathogens showed more specific inhibition patterns

*P. infestans* was the most sensitive pathogen of our study, mostly because it was inhibited by a higher number of strains belonging to the generalist inhibitor clusters mentioned above. In addition, it was also inhibited by strains from two specialist inhibitors clusters, one in the Pseudomonadales order, made of strains all belonging to the *Pseudomonas* genus (Figure 4a), and one in the Propionibacteriales order (filamentous strains), all belonging to the *Nocardioides* genus (Figure 4b).

The presence of the *Pseudomonas* as a specialist inhibitors cluster in our study was surprising because this genus is among the most commonly reported for its biocontrol potential against several pathogens (Anzalone et al., 2021; Berg et al., 2006, 2002; Caulier et al., 2018; Krzyzanowska et al., 2012; Upreti and Thomas, 2015). Their potential as biocontrol agent is widely recognized (Dimkić et al., 2022), and two *Pseudomonas* strains were approved in the EU for biocontrol use (“EU Pesticides Database - Active substances,” 2026).

As for *Nocardioides,* members of this genus are commonly found in soil and among plant endophytes (Han et al., 2013; Qin et al., 2012). They are currently studied for their bioremediation potential (Ma et al., 2023) but to our knowledge, they have not yet been reported as biocontrol organisms, except in our previous study on the same strain collection, where they strongly inhibited Pinf sporangia germination (Pichon et al., 2026).

The presence of many strains showing specific activity against Pinf in this collection is compelling and may be hypothesized to be linked to the fact that strains were isolated from Pinf-challenged plants. However, we consider such an influence unlikely in this case because, although strong shifts were observed in the rhizosphere upon infection with Pinf, the isolated strains corresponding to enriched amplicon sequence variants were overall not more efficient against this pathogen (Pichon et al., 2026). Furthermore, Pinf is frequently reported as the most sensitive pathogen to bacterial inhibition when it is included in screenings with other pathogens, independently of the strains’ origin (Caulier et al., 2018; Guyer et al., 2015; Liu et al., 2017).

Lastly, *Pythium ultimum* had a distinct inhibition profile to all the other pathogens. It was the only pathogen that was not affected by strains from the Bacillales generalist inhibitor cluster (Figure 4a). While it was antagonized by strains from the Streptomycetales cluster, those only represented 53% of the strains active against it, far below the 70% to 80% observed for the other pathogens (Figure 5b). Outside of those, the strains antagonistic to Pult included specialist inhibitors grouped into clusters among the Pseudomonadales (*Pseudomonas* genus) and three orders that displayed hardly any activity against the other pathogens: the Micrococcales (all belonging to the *Frigoribacterium* and *Curtobacterium* genera), Sphingobacteriales (*Pedobacter* genus), and Rhizobiales (*Phyllobacterium* genus). The specialist inhibitors taxa have been so far less studied, but they have nevertheless been reported previously to display antagonistic activity: *Frigoribacterium* strains were reported to inhibit Pinf (Bruisson et al., 2019) and the bee pathogen *Ascosphaera apis* (Khan et al., 2020). *Curtobacterium* has been reported to inhibit the growth *Xanthomonas translucens*, the causal agent of bacterial leaf streak disease (Afkhamifar et al., 2023), and the rice fungal pathogen *Magnaporte oryzae* (Sahu et al., 2021). *Pedobacter* strains inhibited the growth of wheat pathogen *Fusarium graminacearum* (Besset-Manzoni et al., 2019) and of Rsol when co-cultured with a *Pseudomonas* strain (Garbeva and de Boer, 2009). Finally, the inhibitory activity of *Phyllobacterium* strains observed in our study was expected, as this genus included some of the strongest inhibitors of Pinf sporangial germination in our previous screening (Pichon et al., 2026). The absence of inhibition against the same pathogen in the current study may indicate that despite their ability to inhibit sporangia germination, these strains are unable to inhibit vegetative growth. In addition, broad-spectrum fungal inhibition from *Phyllobacterium* strains were also reported (Lambert et al., 1990).

The lack of sensitivity of Pult to Bacillales strains from our collection does not mean that Pult is unaffected by this order more generally. Indeed, *Bacillus* remains among the most frequently reported antagonistic genus against this pathogen (Idris et al., 2008; Lara-Capistran et al., 2020). Pult nevertheless appears to be less susceptible to this taxon than other pathogens. A similar pattern was observed in a screening of 100 strains, in which *Bacillus* strains were inactive against Pult (Beltramino et al., 2025), as well as in a screening of *Bacillus* strains against eight pathogens that also included Pinf and Rsol, where Pult displayed by far the smallest inhibition halos (Liu et al., 2017).

One important difference between the two oomycetes tested in this study is their growth speed, Pult growing substantially faster than Pinf. This parameter likely has a strong impact on the outcome of confrontational assays, as suggested earlier for volatile-mediated interactions (Hunziker et al., 2015). This difference might change when pathogens are growing on their host plant rather than on the culture media we used for the confrontation assays, highlighting the need for ulterior studies in conditions that come closer to the natural situation.

## Conclusion

By screening 600 potato-associated bacterial strains against six major potato pathogens, this study demonstrates the value of multi-pathogen screening for resolving the diversity and specificity of bacterial antagonistic activity. Despite interesting trends, isolation cultivar, plant compartment and cultivation medium had a limited influence on the proportion of antagonistic strains, indicating that potential antagonists were broadly distributed across the sources investigated. In contrast, pathogen identity strongly shaped the observed activity profiles. The bacterial pathogens were highly resistant to inhibition and were almost exclusively affected by broad-spectrum antagonists, whereas the oomycetes displayed more distinct and pathogen-specific inhibition patterns. Pinf was particularly susceptible, while Pult was associated with a distinct set of specialist antagonists.

Antagonistic phenotypes also showed clear phylogenetic patterns, with *Bacillus* and *Streptomyces* containing many broad-spectrum inhibitors, while less commonly studied taxa contained specialist antagonists against the tested oomycetes. These results highlight the benefit of testing bacterial collections against multiple pathogens rather than relying on single-pathogen screens, as this approach can reveal both broad-spectrum candidates and pathogen-specific antagonists.

The screening methods developed here provide a practical and scalable framework for functional characterization of diverse bacterial collections using standard microbiological equipment. Combined with the identification of specialist and broad-spectrum antagonists, this approach provides a useful starting point for the subsequent validation and characterization of candidate mBCAs under conditions that come closer to the real situation, all the way from assays on plant material to greenhouse and ultimately field trials.

## Supporting information

All Supplementary materials and legends

## List of abbreviations

ActIA: Actinomycete Isolation Agar
ARE: Artificial Root Exudate growth medium
Asol: *Alternaria solani*
Dsol: *Dickeya solani*
EU: European Union
MBCAs: Microbial Biological Control Agents
Pcar: *Pectobacterium carotovorum*
PDA: Potato Dextrose Agar Pinf: *Phytophthora infestans*
Pult: *Pythium ultimum*
Rsol: *Rhizoctonia solani*
TSA: Tryptic Soy Agar

## Clinical trial

Not applicable.

## Clinical Trial Number

Not applicable.

## Consent for publication

Not applicable.

## Data Availability

All data and R scripts generated during this study are published on GitHub at https://github.com/VivienPichon/Collection_antagonistic_activity_against_6_pathogens, and on Zenodo at the DOI: 10.5281/zenodo.22258863

## CRediT authorship contribution statement

**Vivien Pichon**: Conceptualization, Methodology, Validation, Formal Analysis, Data Curation, Writing – Original Draft, Visualization, Investigation. **Alisson Gillon**: Methodology, Validation, Investigation, Data Curation, Writing – Original Draft. **Gaëlle Ruffieux**: Validation, Investigation, Writing – Review and Editing. **Tom Luethi**: Validation, Investigation, Writing – Review and Editing. **Camila Morales Undurraga**: Validation, Investigation, Writing – Review and Editing. **Mout De Vrieze**: Methodology, Writing – Review and Editing, Supervision. **Floriane L’Haridon**: Resources, Writing – Review and Editing. **Ola Abdelrahman**: Methodology, Writing – Review and Editing, Supervision. **Laure Weisskopf**: Conceptualization, Methodology, Writing – Original Draft, Supervision, Project Administration, Funding Acquisition.

## Ethics approval and consent to participate

Not applicable.

## Acknowledgements

Financial support from the Gerbert Rüf Stiftung and from the Swiss National Science Foundation (grants 207917 and 229296 to L.W.) is gratefully acknowledged.

