## Supplementary material for "A scalable screening approach reveals phylogenetic patterns of bacterial antagonism against potato pathogens": All Supplementary materials and legends

September 2026

**Figure S1**: Examples of confrontations for each score and each pathogen.

| | | ***Dickeya/Pecto*** | ***Alternaria*** | ***Rhizoctonia*** | ***Phytophthora*** | ***Pythium*** |
|---|---|---|---|---|---|---|
| 0 | No Inhibition. | 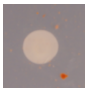 | 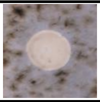 | 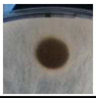 | 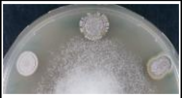 | 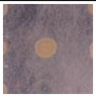 |
| 1 | The pathogen stops at the edge of the colony. | 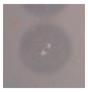 | 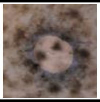 | 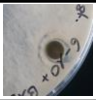 | 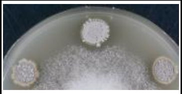 | 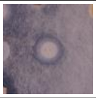 |
| 2 | The pathogen stops between 1 mm and 5 mm away from the colony. | 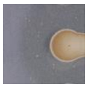 | 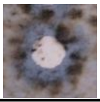 | 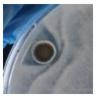 | 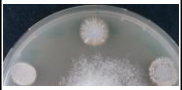 | 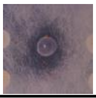 |
| 3 | The pathogen stops more than 5 mm away from the colony but not completely the neighbouring colonies. | 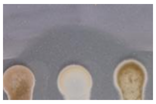 | 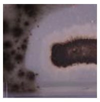 | 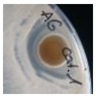 | 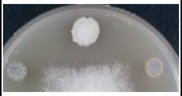 | 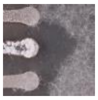 |
| 4 | The inhibition area includes entirely neighbouring colonies. | ⊘ | 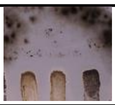 | ⊘ | 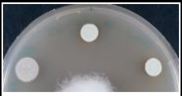 | ⊘ |

**Figure S2**: Distribution of the maximum antagonistic activity of bacterial isolates according to cultivar of origin **(a)**, plant compartment of origin **(b)**, growth medium used for isolation **(c)**. Each subplot is split by strain type (non-filamentous or filamentous) and pathogen. Stacked bar plots show the proportion of strains assigned to each activity level (inactive, low, moderate, high, and outstanding), with each bar representing 100% of the strains within a given category.

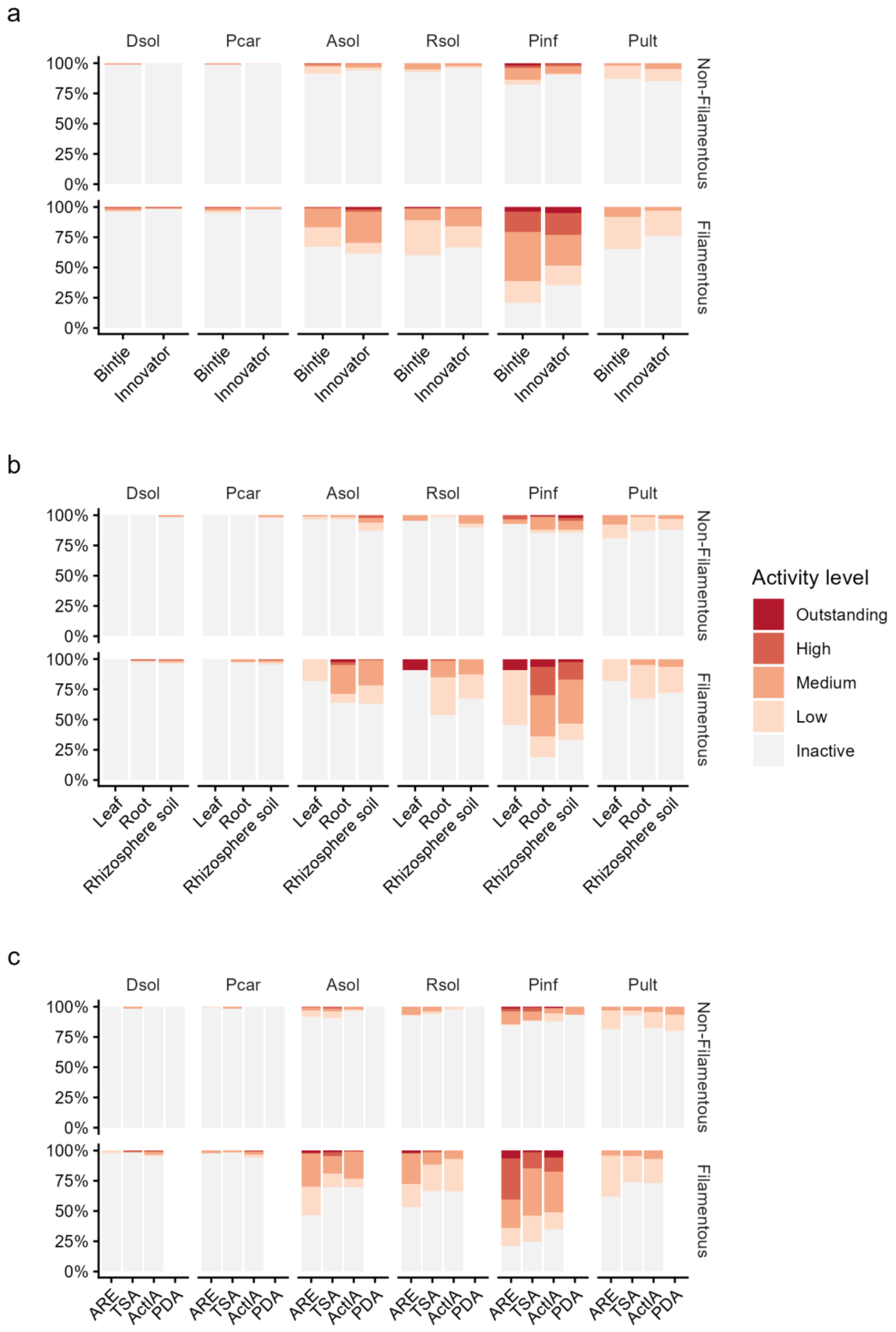

**Figure S3**: Phylogenetic trees showing the antagonistic activity of every bacterial isolate against at least one pathogen for non-filamentous (a) and filamentous (b) strains. Branch colours indicate the order each strain belongs to. The circular heatmap around the trees represents the antagonistic activity level of each strain against each pathogen. From the innermost to the outermost ring, activity is shown against *Dickeya solani* (Dsol), *Pectobacterium carotovorum* (Pcar), *Alternaria solani* (Asol), *Rhizoctonia solani* (Rsol), *Phytophthora infestans* (Pinf), and *Pythium ultimum* (Pult). White colour represents missing data.

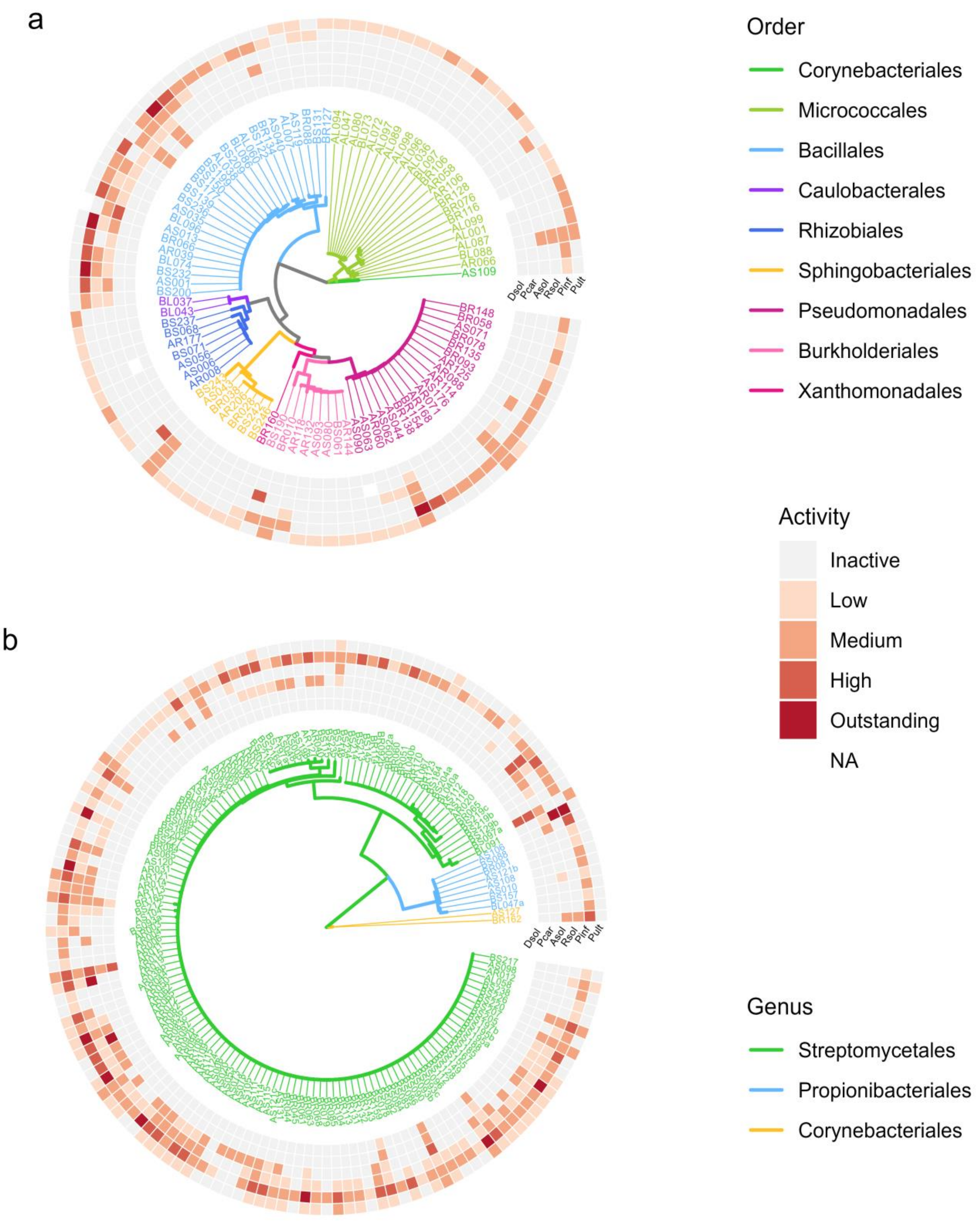

**Figure S4**: Distribution of the antagonistic activity levels of the predominant orders of the bacterial collection against Dsol **(a)**, Pcar **(b)**, Asol **(c)**, Rsol **(d)**, Pinf **(e)** and Pult **(f)**. Predominant orders were defined as those which included at least 10 strains in the collection, and exhibiting at least 15% of active strains. Stacked bar plots show the proportion of strains assigned to each activity level (inactive, low, moderate, high, and outstanding), with each bar representing 100% of the strains within a given category.

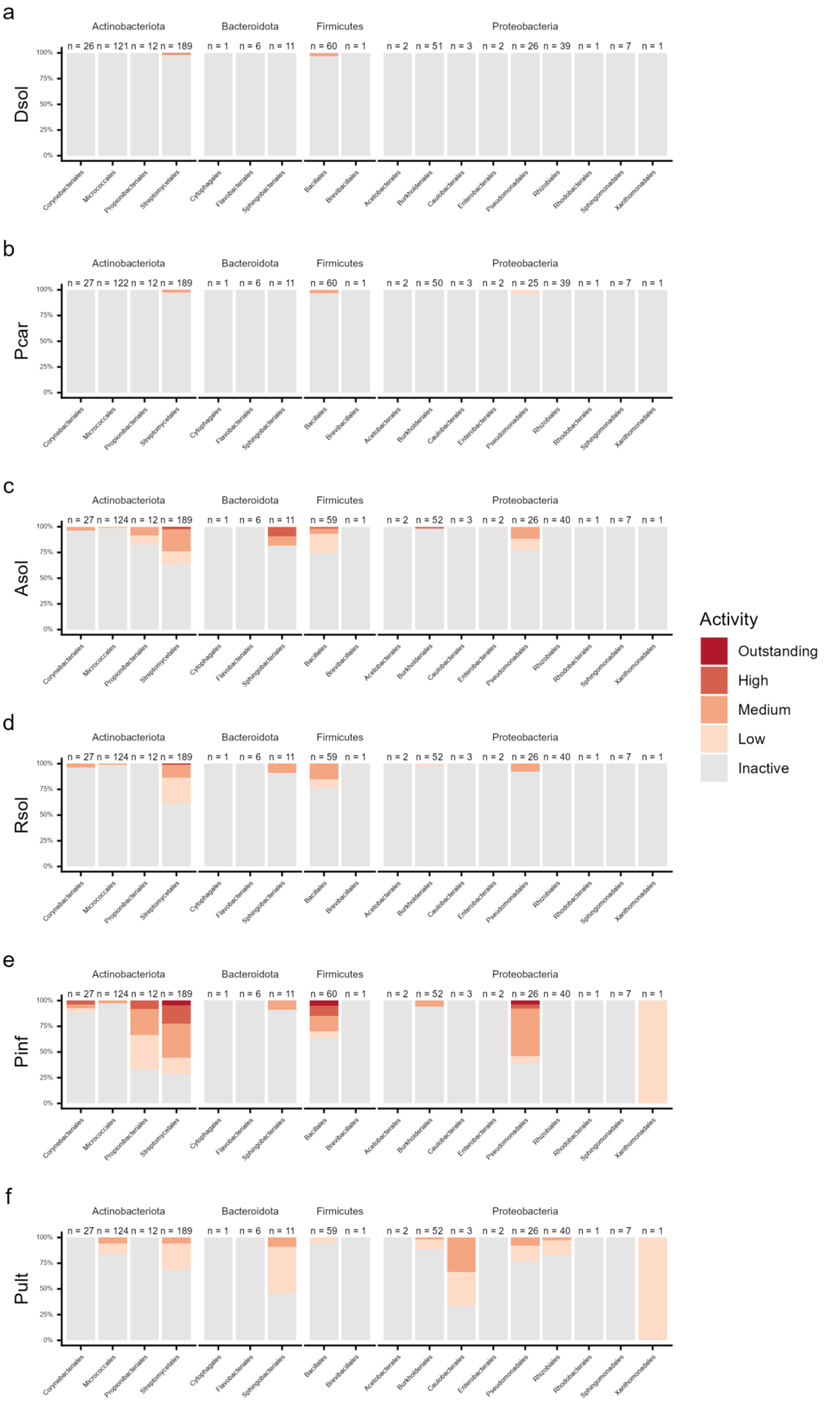
